# HIV-1 Induction of Tolerogenic DCs is Mediated by Cellular Interaction with Suppressive T Cells

**DOI:** 10.1101/2021.09.01.458353

**Authors:** Cecilia Svanberg, Sofia Nyström, Melissa Govender, Pradyot Bhattacharya, Karlhans F Che, Rada Ellegård, Esaki M Shankar, Marie Larsson

## Abstract

HIV-1 infection gives rise to a multilayered immune impairment in most infected individuals. The crosstalk between Dendritic cells and T cells plays an important part in the induction of immune responses. The chronic presence of human immunodeficiency virus (HIV)-1 during the dendritic cells (DCs) priming and activation of T cells promotes the expansion of suppressor cells in a contact dependent manner. The mechanism behind the T cell side of this HIV induced impairment is well studied, whereas little is known about the reverse effects exerted on the DCs in this setting.

Here we assessed the phenotype and transcriptome profile of mature DCs that have been in contact with suppressive T cells. The DCs in the HIV exposed DC-T cell coculture obtained a more tolerogenic/suppressive phenotype with increased expression of e.g., PDL1, Gal-9, HVEM, and B7H3, mediated by their cellular interaction with T cells. The transcriptomic analysis showed a clear type I IFN response profile as well as an activation of pathways involved in T cell exhaustion.

Taken together, our data indicate that the prolonged and strong IFN type I signaling induced by the presence of HIV during DC-T cell cross talk could play an important role in the induction of the tolerogenic DCs and suppressed immune response.

## Introduction

Dendritic cells (DCs) serve as a first line of defense against many microbes, including human immunodeficiency virus-1 (HIV) (Steinman 2007, Chougnet and Gessani 2006), and their ablation has been linked to development of autoimmunity (Ohnmacht et al. 2009). Immature DCs become activated upon engagement of their pattern recognition receptors (PRRs) with pathogen-associated molecular patterns (PAMPs) (Steinman 2007, Lundberg et al. 2014). Following maturation, DCs gain T cell priming abilities owing to increased antigen processing and upregulation of MHC class I and II molecules, co-stimulatory molecules and cytokines that further program the ensuing immune responses. The interaction of DCs with a wide range of cells, such as natural killer (NK) cells, NKT cells, basophils, regulatory T cells (Tregs), memory T cells, epithelial cells, and stromal cells has a huge impact on the ensuing immunological outcome (Pulendran, Tang and Manicassamy 2010, Ellegard et al. 2018, Tong et al. 2021). The interaction between immune cells and the network of signaling cascades are complex, and it will be the sum of the signaling by co-stimulatory and co-inhibitory molecules as well as the type of antigen that determines the fate of the response. For instance, the polarization of naïve T helper (TH) cells occurs via interdependent routes resulting from the microenvironment created by DC-specific stimulation and accessory cell interactions, culminating in TH-subset differentiation and regulatory T cell responses (Pulendran et al. 2010, Luckheeram et al. 2012, Basu et al. 2021, Yin, Chen and Eisenbarth 2021).

The prolonged response to chronic antigenic stimulation such as chronic viral infection can lead to exhausted T cells with lost effector functions or give rise to regulatory/suppressive T cells (Dowling et al. 2018, Blank et al. 2019, Im and Ha 2020). Induced regulatory/suppressive T cells are implicated with the control of inflammatory diseases (Lan et al. 2012, Saxena et al. 2021, Abdel-Gadir, Massoud and Chatila 2018, Sakaguchi et al. 2008). The subtypes of inducible regulatory T cells are complex and consisting of many different subgroups (Mason et al. 2015). The properties of the regulatory/suppressive cells activated depend on the microenvironment in which they were created/activated and signals they received, for instance, while prolonged IL-10 exposure can generate one subtype, exposure to transforming growth factor-β (TGF-β) can result in another (Levings et al. 2002, Chen et al. 2003, Iberg and Hawiger 2020, Roncarolo et al. 2018). Regulatory/suppressive T cells can express several co-inhibitory molecules such as programmed cell death 1 (PD1), cytotoxic T lymphocyte antigen 4 (CTLA-4), lymphocyte activation gene 3 (LAG-3), T cell immunoglobulin and mucin domain-containing protein 3 (TIM3), CD160 and, B lymphocyte-induced maturation protein-1 (BLIMP-1), and depending on the frequency of expression of these markers, the T cells exhibit different levels of suppression (Fromentin et al. 2016, Du et al. 2020).

The presence of impaired/regulatory or tolerogenic DCs have been described in different settings such as tumor microenvironment and chronic infections (Shurin, Ma and Shurin 2013, Schmidt, Nino-Castro and Schultze 2012). Tolerogenic DCs can suppress T cell responses and induce differentiation of regulatory/suppressive T cells (Kushwah and Hu 2011). These DCs can have elevated levels of IL-10, TGF-β, indoleamine 2,3 dioxygenase (IDO), cyclooxygenase-2 (COX-2), retinoic acid (Kushwah and Hu 2011), and expression of co-inhibitory molecules, notably programmed death-ligand (PD-L) 1/PD-L2 (Riley 2009), B7H3/B7H4 (Loos et al. 2010), HVEM (del Rio et al. 2010), and/or galectin-9 (Gal-9) (Lu et al. 2019). It’s been difficult to define a specific phenotype and level of inhibition of T cells, which demonstrate the plasticity of DCs, i.e., ability to adopt to the environmental cues. It has been indicated that during chronic conditions such as viral persistence, the type I interferons (IFNs) can have immunosuppressive effects and program individual immune cells such as DCs into cells with more immunosuppressive functions (Cunningham et al. 2016). Furthermore, direct exposure to HIV gp120 could impair the functional attributes of DCs (Chougnet and Gessani 2006, Knight and Patterson 1997, Izquierdo-Useros et al. 2014, Quaranta et al. 2006, Shan et al. 2007).

DC-signaling via co-stimulatory molecules, such as CD80, CD86, and CD40, or co-inhibitory molecules such as PDL1 during the DC-T cell interactions shapes the T cell response. This engenders in enhanced bidirectional cell survival, by triggering T cell activation, which in turn activates the DCs to secrete IL-12 that polarizes the immune response towards the TH1 (Kelsall et al. 1996, Hirohata 1999, Wesa and Galy 2002, Tay et al. 2017). We have previously reported the effect HIV exerts on the DC-T cell crosstalk during priming and activation using an in vitro model and have shown that HIV-exposed mature DCs induce T cells with immune suppressive properties (Che et al. 2010, Che et al. 2012, Shankar et al. 2011). In those studies the T cells had increased expression of several co-inhibitory molecules, including PD1, CTLA4 and LAG-3 and CD160, (Che et al. 2010, Che et al. 2012, Shankar et al. 2011).

In this study, we have explored the effects on the DC side in the DC-T cell coculture exposed to HIV to elucidate how the suppressive T cells found in this setting affects the DCs. This has not, as far as we are aware, been studied in the setting of HIV. To this end we have characterized DC phenotype by flow cytometry and by transcriptomics. We found that interaction with T cells in the presence of HIV, but not HIV alone, induced DCs with tolerogenic properties characterized by increased expression of negative co-stimulatory molecules such as PDL1, GAL-9, and herpes virus entry mediator (HVEM). At the transcriptome level we did found a sustained type I IFN response and downstream type I IFN induced pathways in HIV-exposed DCs from DC-T cell cocultures that may explain the induction of tolerogenic properties of the DCs after T cell interaction.

## Materials and methods

### Virus

HIV-1_BaL_/SUPT1-CCR5 CL.30 was produced using chronically infected cultures of ACVP/BCP Cell line (No. 204), originally derived by infecting SUPT1-CCR5 CL.30 cells (graciously provided by Dr. J. Hoxie, University of Pennsylvania) with an infectious stock of HIV-1_BaL_ (NIH AIDS Research and Reference Reagent Program, Cat. No. 416, Lot 59155). Virus was purified by continuous flow centrifugation using a Beckman CF32Ti rotor at 30,000 rpm (∼90000xg) at a flow rate of 6 liters/hour followed by banding for 30 minutes after sample loading. Sucrose density-gradient fractions were collected, virus-containing fractions pooled and diluted to <20% sucrose, and virus pelleted at ∼100,000xg for 1 hour. The virus pellet was resuspended at a concentration of 1,000xg relative to the cell culture filtrate and aliquots frozen in liquid N_2_ vapor.

### Reagents

RPMI1640 was supplemented with 10mM HEPES, 20µg/mL gentamicin (Fisher Scientific, Leicestershire, UK), 2mM L-glutamine (Sigma-Aldrich, St. Louis, MO), and 1% plasma or 5% heat-inactivated pooled human serum (5% PHS). Recombinant human granulocyte-macrophage-colony stimulating factor (rhGM-CSF) (100IU/mL) (Preprotech, London, UK) and recombinant human interleukin-4 (rhIL-4) (300U/mL) (Preprotech, London, UK), were used for the in vitro differentiation of DCs.

### Propagation and maturation of monocyte derived Dendritic cells

Buffy coats or whole blood were obtained from healthy individuals (ethical permit M173-07) and peripheral blood mononuclear cells (PBMCs) were isolated by density-gradient centrifugation over Ficoll-Paque™ (Amersham Pharmacia, Piscataway, NJ). Monocytes were selected by plastic adherence after incubation of PBMCs in tissue culture dishes (BD Falcon, Franklin Lakes, NJ) for one hour. The plates were then washed with RPMI1640 to remove non-adherent cells. The remaining adherent cells were cultured in 1% single donor plasma medium supplemented with rhGM-CSF (100 U/mL) and rhIL-4 (300 U/mL) and incubated in a 5% CO_2_ environment at 37°C. The rhGM-CSF and rhIL-4 cytokine supplement was replenished every second day to facilitate differentiation of CD14+ progenitor cells into immature DCs, and the cells were harvested on day 5. The purity and readiness of immature DCs was assessed by flow cytometry on a FACSCanto II (BD Immunocytometry Systems, San Jose, CA). The DCs were phenotyped using phycoerythrin-conjugated (PE) monoclonal antibodies (mAbs) against CD83 and CD14 setting the cutoff for positive cells with the isotype control IgG_2a_ (BD Pharmingen, Franklin Lakes, NJ). Immature DCs were transferred to new plates at a concentration of 4×10^5^cells/mL, and maturation was induced by adding 30ng/ml polyinosinic acid: polycytidylic acid (Poly I:C) (Sigma-Aldrich, St. Louis, MO). The maturation of DCs was assessed, after 24h of incubation, by CD83 surface expression by flow cytometry as described above (Supplementary Figure 1).

### HIV exposure of mature DCs

Following maturation, HIV-1_BaL_ (Lot No P4143, 4235, 4238, 4213: 750 ng/mL p24 equivalents/mL corresponding to ∼2 MOI, a concentration reported to occur in vivo, was added to the DCs and the cells were incubated for an additional 24 hours. The unbound viruses were washed off the DCs by rinsing the plates with RPMI and changing the media before use in the assays. HIV unexposed mock mature DCs served as controls, and DC viability following HIV exposure was examined by trypan blue exclusion method.

### Setup of the DC-T cell coculture

Cocultures of HIV-exposed and mock-exposed mature DCs with naïve T cells were set in 96-well flat-bottom cell culture plates. The naïve T cells were isolated from allogenic non-adherent cells by a negative selection using magnetic beads coupled with anti-CD56, anti-CD19, anti-CD45RO and anti-CD14 magnetically tagged antibodies (MACS, Miltenyi Biotec, Auburn, CA) to deplete NK cells, B cells, memory T cells, and monocytes, respectively. The naïve T cell preparation was cocultured with mock or HIV-pulsed DCs at a 1:10 ratio and incubated at 37^º^C in a 5% CO_2_ incubator. An aliquot of the DCs was stored frozen at -80°C in FBS containing 8% DMSO and used for restimulating. The cocultures were restimulated on day 7 with the mock or HIV treated DCs from the same donor. On day 8, the cocultures were harvested.

### T cell proliferation assay

After restimulation, the cocultures were pulsed with 2µCi/µl of [^3^H] labelled thymidine and incubated for about 16 h before harvesting. The radioactive thymidine incorporated into the DNA of proliferating T cells was determined by liquid scintillation counting using a micro-β counter (PerkinElmer, Waltham, MA, USA).

### Immunostaining of DCs and T cells

Direct conjugated mAbs against CD3-PE-Cy7, CD8–PE, CD4–APC, CD1c-BV421, HLA-DR-PERCP, B7H3-APC, B7H1-PE, NOS1-FITC, PDL2-APC, HVEM-PE, Arginase 1-PerCP-eFluor 710, IDO-FITC, CD80-APC, CD86-PE, CD85d (ILT4)-PerCP-eFluor 710, COX2-FITC, CD30L/TNFSF8-APC, Galectin-9-PE, and B7-H4-Alexa Fluor 700 were used for DC and T cell phenotyping before and after the DC–T cell interaction in the coculture. Data was acquired using FACS canto II and the FlowJo software (Treestar, OR, USA) were used for data analysis.

### Separation of DCs from the DC-T-cell coculture and RNA sequencing

Two kits of magnetically labelled beads were used to separate DCs from T cells after coculture. Beads for removal of dead cells (Miltenyi) and beads targeting CD1c+ cells (Miltenyi) for positive selection of DCs and consequently negative selection of the T-cells. Thereafter cells were lysed, and RNA was extracted using Isolate II RNA Mini or Micro Kit (Bioline, UK). An amount of 5 ng of total RNA per sample was used for transcriptome amplification using NuGEN’s Ovation RNA-Seq V2 kit (San Carlos, CA, USA). Library quality and size distribution was determined using Agilent Bioanalyzer 2100. A total of 16 libraries from 8 different donors (two conditions/donor) were prepared and ran in two different runs on the Illumina NextSeq500 platform (San Diego, CA, USA). FASTQ files were analyzed using UPPMAX, and quality was checked using the fastQC and multiQC programs. After quality controls libraries were trimmed using trimmomatic followed by mapping of the libraries using STAR to the human reference genome hg19. FeatureCounts was applied for retrieving counts for each mapped gene. Using R/DeSeq2 data was normalized and differentially expressed genes were determined. Thereafter the data was visualized using Ingenuity Pathway Analysis (IPA, Qiagen), R analysis and Gene Ontology (GO) Enrichment Analysis (Geneontology.org). A p-value cut-off of 0.05 was set as significant for affected molecules/pathways.

### Quantitative Polymerase Chain Reaction

Cryopreserved cell lysates of DCs (24h exposure), enriched DCs or T-cells from cocultures and whole cocultures were used for RNA extraction, cDNA synthesis and quantitative (q) PCR analysis. RNA was extracted and first strand cDNA was synthesized using the superscript III RT kit (Invitrogen^™^, USA). The primers used for qPCR were retrieved from the primer bank found at http://pga.mgh.harvard.edu/primerbank/. Expression of negative co-stimulatory molecules together with β-actin and GAPDH endogenous controls were analyzed using CFX96 Touch Real-Time system (BIO-RAD Inc.) in 96-well MicroAmp^®^ optical fast-plates (Applied Biosystems^®^) and Fast SYBR Green^®^ Master Mix (Applied Biosystems^®^). Results were firstly normalized according to the 2^-ΔΔCt^ method, where *Ct* refers to the threshold value, and thereafter further transposed and normalized as fold-change to Mock.

### Statistical Analysis

All data were analyzed using MS Excel and GraphPad Prism 9 software (GraphPad, La Jolla, CA, USA) using paired t test to compare between the two groups. Data obtained from qPCR experiments were normalized, each data set was divided by the sum of all values in the data set and each value within a group is presented as percentage. For all tests, a *P* value of <0.05 was considered significant, and the measure of significance was represented by \**P<*0.05, \*\**P<*0.005 and \*\*\**P<*0.001.

## Results

### Presence of HIV during DCs priming of naïve T cells leads to suppressive T cell responses

In an HIV infected individual there will be virions in the lymphoid tissue, affecting the priming and reactivation of T cells and the quality of the immune responses (Boasso, Shearer and Chougnet 2009, Estes 2013, Pantaleo et al. 1991, Nguyen et al. 2020). Here we tried to mimic the in vivo setting in the lymph nodes, by using an experimental allogenic HIV priming model with mature DCs and naïve T cells **(Figure 1)** where we assessed the effects the DC-T cell crosstalk exerts on the DCs’ phenotype and transcriptome profile. Mock or HIV exposed mature DCs were cocultured with CD3^+^CD45RA^+^ bulk naïve T cells. On day seven, cocultures were restimulated with the same donor DCs added on day 1 and T cell proliferation was measured by [^3^H] thymidine incorporation. HIV exposure of mature DCs resulted in impaired T cell priming compared to the mock treated mature DCs illustrated by decreased proliferation **(Figure 2A)**. DCs exposed to HIV before the coculture with T cells had the same viability and phenotype as the mock-treated cells **(Supplementary Figure 1)**. There was a low level of HIV gag transcripts in cocultures of HIV exposed DCs **(Figure 2B)**, which confirms our previous data that there were a minimal level of HIV replication (Che et al. 2010) in the system.

**Figure 1.**
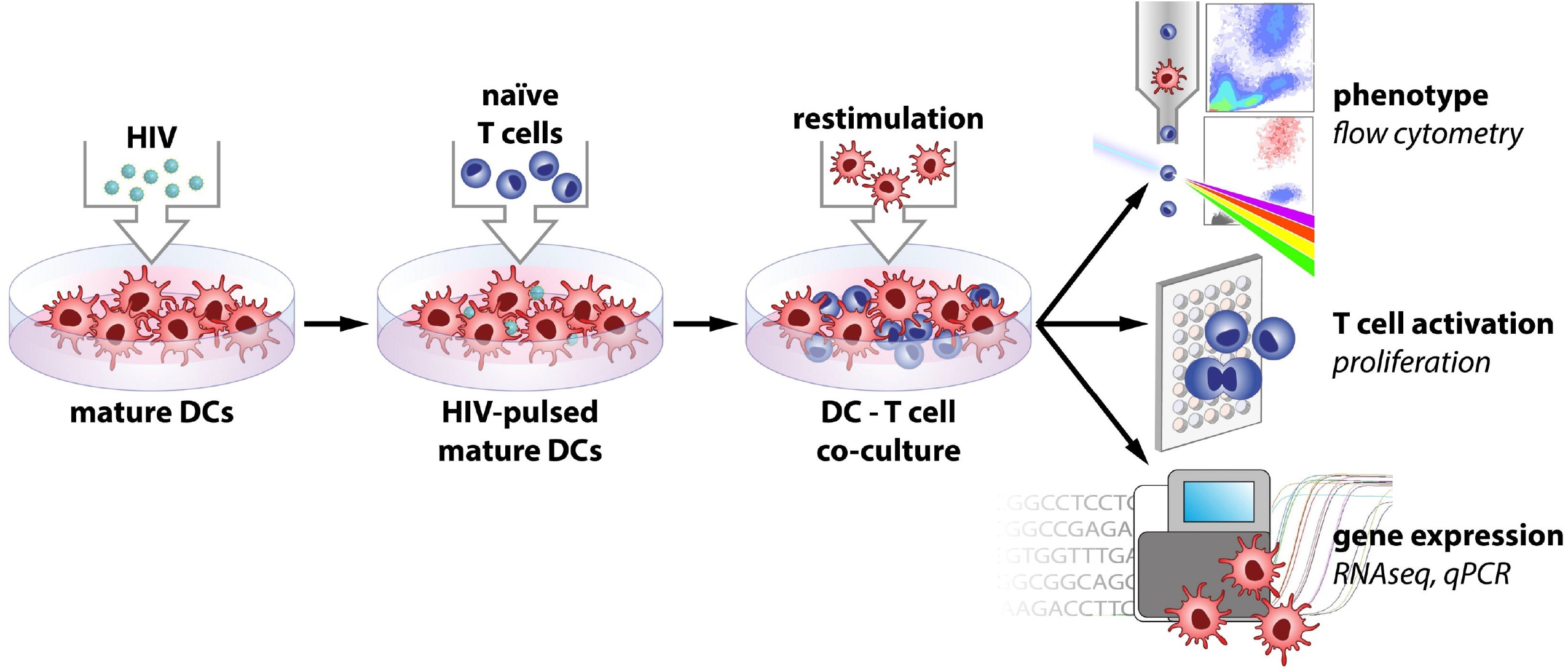
Model of experimental setup.

**Figure 2.**
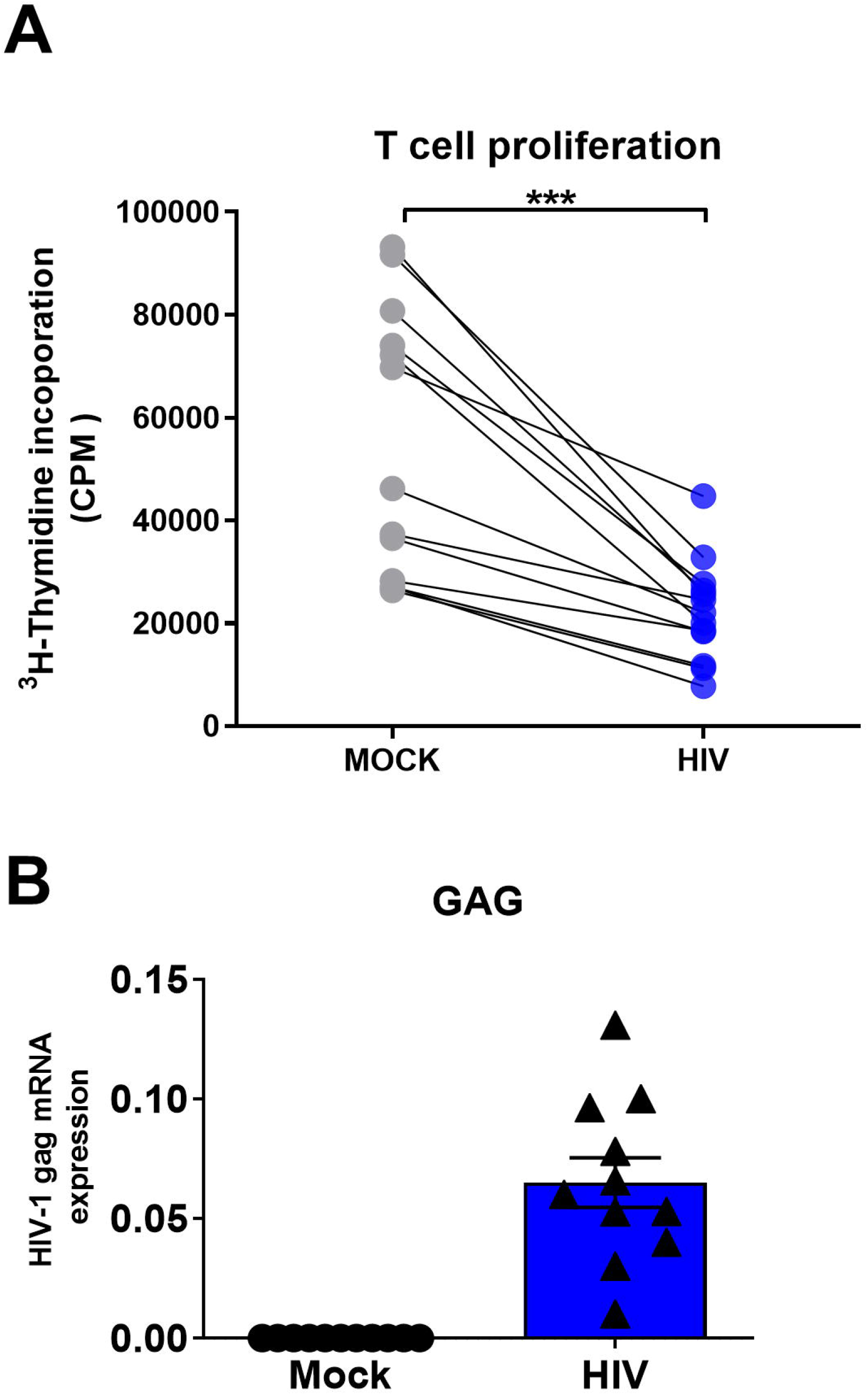
Presence of infectious HIV impairs the ability of DCs to prime naïve T cell responses. Mature DCs were pulsed with infectious HIV-1_BaL_ (CCR5-tropic) overnight, washed twice, and cocultured with naïve bulk T cells at a ratio of 1:10 (10^4^ DCs and 10^5^ T cells). The priming cultures were restimulated with 10000 DCs per well after 7 days of coculture, and [^3^H] thymidine incorporated into the DNA of proliferating T cells was determined on day 8 by liquid scintillation counting using a micro-β counter. Values expressed as counts per minute (CPM). **(A)** HIV effects on the ability of DCs to induce T-cell proliferation <40000 CPM and **(B)** HIV effects on the ability of DCs to induce T cell proliferation >40000 CPM. N=11-13. Normalized values for assays performed with CCR5-tropic HIV-1_BaL_ (<40000, n=10; >40000, n=6). ^***^*P*<0.001, unpaired t-test.

### HIV Exposure Leads to Differentiation of mature DCs to Tolerogenic DCs Following Cellular Crosstalk with T Cells

Today it is clear that the DCs phenotype and function can be further modified by the environmental stimulus and cellular interactions (van den Biggelaar et al. 2020, Miller and Bhardwaj 2013, Wu and KewalRamani 2006, Rodriguez-Plata et al. 2012). We investigated the effect of cellular interaction with T cells on mature DCs exposed to HIV, with focus on established co-inhibitory factors known to be expressed by DCs with a tolerogenic phenotype. We exposed matured DCs to mock, or HIV-1_BaL_ for 24h and found that the HIV exposure alone, had no effect on DCs’ gene expression levels of PDL1, decoy receptor (DcR) 2, Gal-9, HVEM, tryptophan 2,3-dioxygenase (TDO), or cyclooxygenase (COX)-2 **(Figures 3A-B)**. The only factor that increased in HIV exposed DCs compared to mock treated DCs, was IDO **(Figure 3B)**. We further investigated the expression of co-inhibitory factors and enzymes, in the mock vs HIV DC-T cell cocultures of mock or HIV treated DCs after restimulation. We found increased gene expression levels of Gal-9, PDL1, and DcR2 **(Figure 3C)**, as well as IDO, TDO, and COX-2 in restimulated cocultures with DCs exposed to HIV **(Figure 3D)**. PDL1 and IDO levels were already elevated at 2 and 5h after restimulation of the coculture with HIV exposed DCs **(Supplementary Figure 2)**. We next confirmed the RNA expression findings at the protein level by flow cytometry analysis of the DCs in the DC-T cell cocultures after 24h of restimulation **(Figure 4A-B)**. We found significantly increased levels of PDL1, HVEM, Gal-9, B7H3 (CD276), PDL2, CD85, CD30L, and DR4 in HIV exposed DCs compared to mock in restimulated cocultures (**Figure 4A-B**). HIV exposure did not alter the protein expression of IDO, COX-2, NOS1, and arginase 1 **(Figure 4B)**. Taken together, this comprehensive analysis suggests that the crosstalk between DCs and T cells in the presence of HIV induces DCs with a tolerogenic phenotype **(Figure 4C)**.

**Figure 3.**
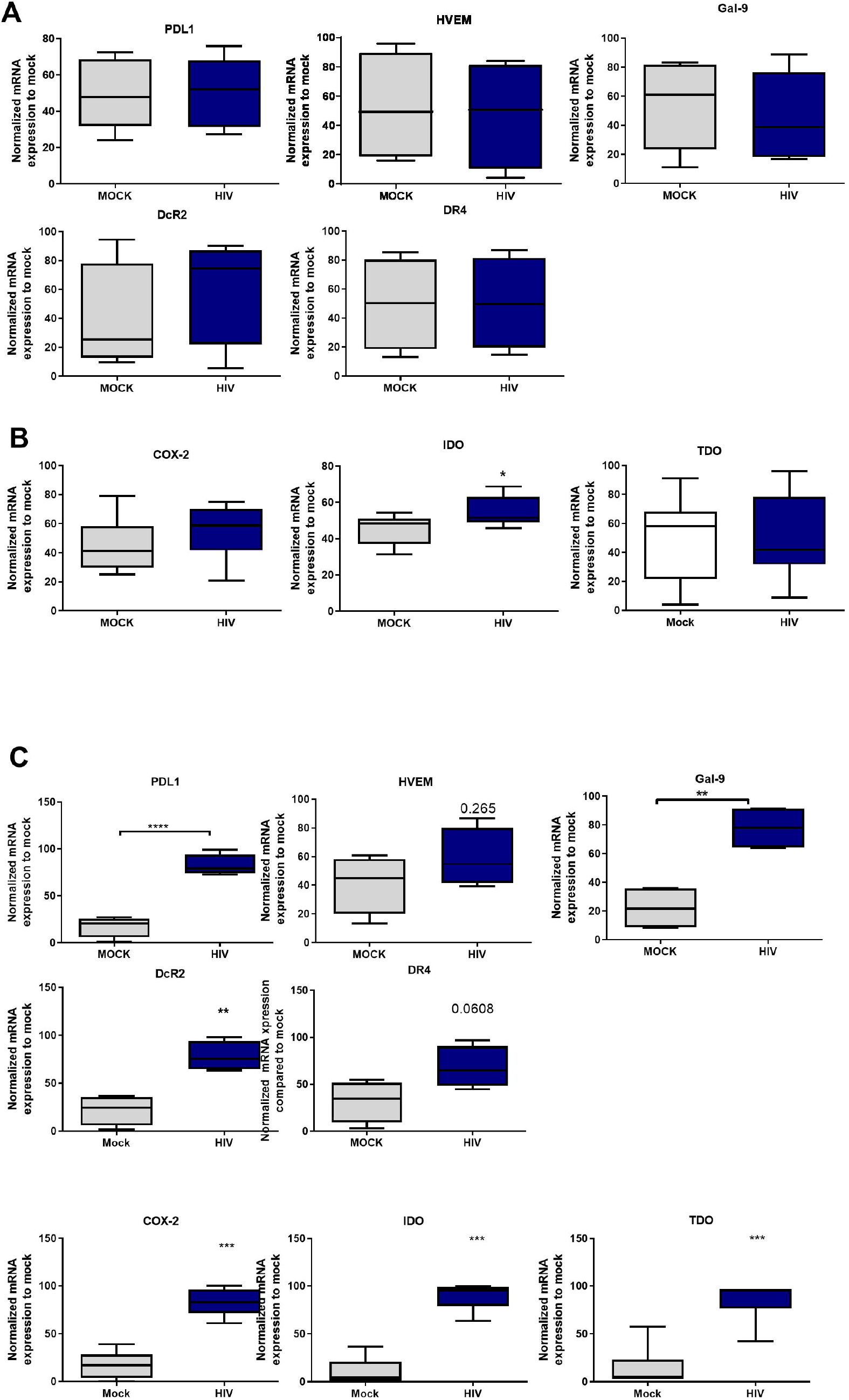
Presence of HIV in DC-T cells coculture give rise to DCs with an increased gene expression of factors associated with a tolerogenic phenotype. Mature DCs were pulsed with mock and HIV-1_BaL_ (HIV), and cultured 24h, washed and studied for gene expression levels of **(A)** PDL1, HVEM, Gal-9, DcR2 and DR4 and **(B)** COX-2, IDO and TDO by DCs. N=7. DC-T cell cocultures, with or without HIV were harvested after 8 days. The gene expression levels in the day 8 coculture of **(C)** PDL1, HVEM, Gal-9, DcR2, DR4 **(D)** IDO, COX-2, and TDO examined by PCR (N=5-11). The experiments were normalized to mock, and statistics performed. ^*^*P*<0.05, ^**^*P*<0.005, ^***^*P*<0.001, unpaired t-test.

**Figure 4.**
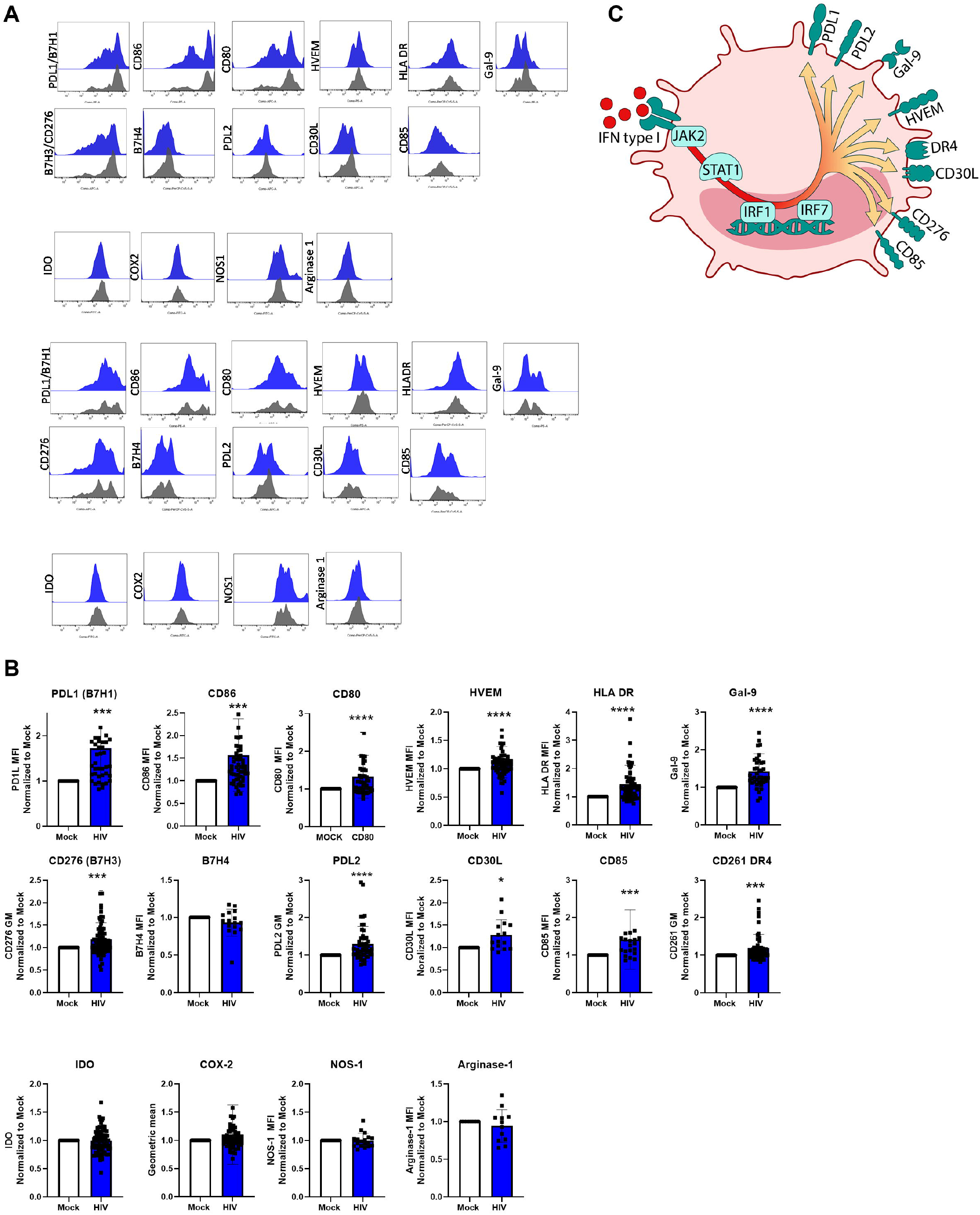
Presence of HIV in DC-T cells coculture give rise to DCs with upregulated protein expression of molecules associated with a tolerogenic phenotype. DC-T cell cocultures, with or without HIV were harvested after 8 days. The DCs in the coculture were identified by expression of CD1c. DC expression-levels of PDL1, CD80, CD86, HVEM, HLA-DR, Gal-9, B7H3, B7H4, PDL2, CD30L, CD85, IDO, COX-2, NOS1, and arginase 1 were examined. Representative histograms of 2 donors show the expression of these markers on CD1c positive DCs **(A)**. Graphs of normalized MFI values of the markers on CD1c positive DCs **(B)**. Illustration of the tolerogenic DC **(C)**. The experiments (N=12-60) were normalized with mock set as one, and statistics performed. ^*^*P*<0.05, ^**^*P*<0.005, ^***^*P*<0.001, unpaired t-test.

### The transcriptomic profile showed type I IFN pathway activation in the DCs from HIV conditioned DC-T cell cocultures

To assess the transcriptomic profile of the DCs in the DC-T cell coculture we sorted the DCs after 24h restimulation and performed RNA sequencing (see model figure 1A). Partial least squares discriminant analysis (PLS-DA) model was used to assess if there were underlying relationships between the DCs from the mock and HIV DC-T cell cocultures. The PLS-DA revealed a clear separation between the DCs samples from the mock compared to the DC samples from the HIV exposed DC-T cell coculture (**Figure 5A**). To further examine the transcriptomic data, we used IPA to analyze the genes and pathways that separated the DCs sorted from the HIV coculture vs mock coculture. The graphic summary/overview of the main pathways affected clearly illustrated a strong activation of type I IFN response, an upregulation of innate immune factors and pathways associated with inhibition of viral replication (**Figure 5B**). The T cell exhaustion signaling pathway and Interferon signaling were among the top pathways with positive Z-scores 1.4 and 3 respectively (**Table 1**). The canonical pathways that were inhibited/decreased included the ICOS-ICOSL Signaling in T helper cells, and TGF-β Signaling, and PKC□ Signaling in T lymphocytes (**Table 2**).

**Figure 5.**
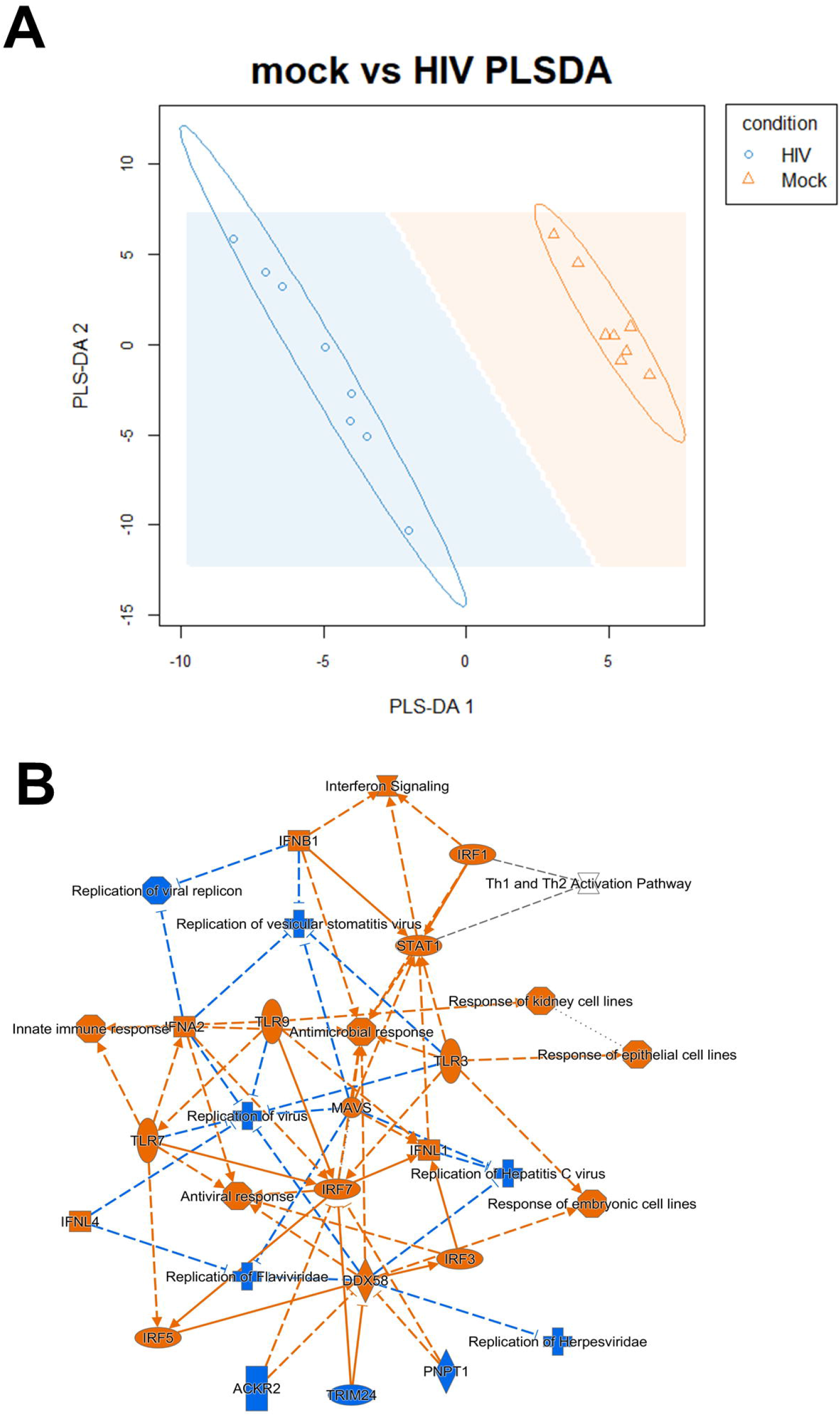
Transcriptomic data shows a high separation of the mock vs HIV DC from the DC-T cell coculture and clear type I IFN signaling profile in HIV exposed DC cocultures. PLS-DA analysis was performed to model relationship between HIV exposed and mock treated DCs after coculture with T cells (N=8) **(A)**. Transcriptomic data set including 8 individual donors/experiments were analyzed using Ingenuity Pathway Analysis (IPA). IPA selected pathways were visualized as network with a threshold for p-values set to -log 1.3 (p<0.05) and presented as positive activation Z-score in orange and negative in blue **(B)**.

**Table 1:**
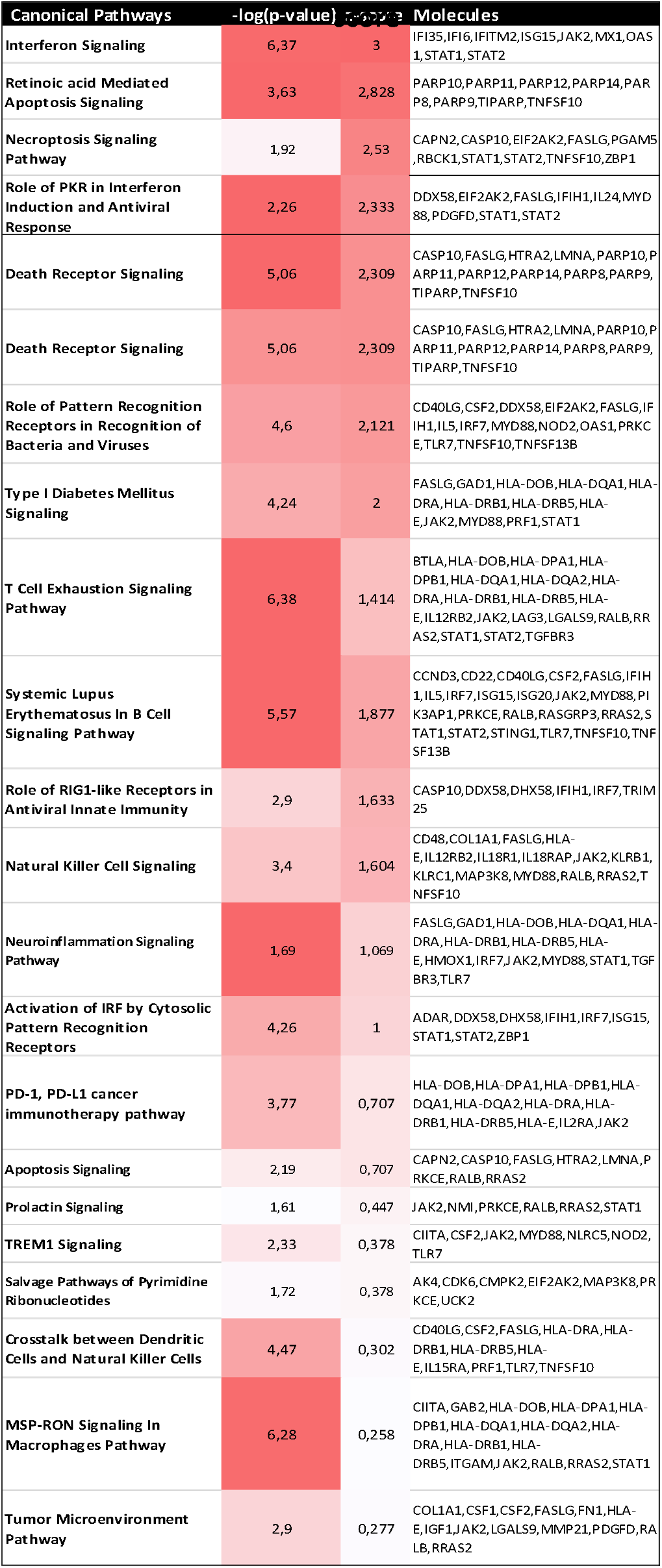
Canonical pathways with positive Z-

**Table 2:** Canonical pathways with negative Z-score

| Canonical Pathways | -log(p-value) | z-score | Molecules |
| --- | --- | --- | --- |
| iCOS-iCOSL Signaling in T Helper Cells | 2,99 | -2,333 | CD4, CD40LG, GAB2, HLA-DOB, HLA-DQA1, HLA-DRA, HLA-DRB1, HLA-DRB5, IL2RA, PLEKHA1 |
| TGF-β Signaling | 1,3 | -2,236 | IRF7, RALB, RRAS2, RUNX2, SMAD9, VDR |
| PKCθ Signaling in T Lymphocytes | 1,54 | -1,667 | CD4, HLA-DOB, HLA-DQA1, HLA-DRA, HLA-DRB1, HLA-DRB5, MAP3K8, RALB, RRAS2 |
| SPINK1 General Cancer Pathway | 1,92 | -1,633 | JAK2, MT1E, MT1F, MT2A, RALB, RRAS2 |
| Systemic Lupus Erythematosus In T Cell Signaling Pathway | 1,62 | -1,5 | CASP10, CASP4, CD40LG, ELF1, FASLG, HLA-DOB, HLA-DPA1, HLA-DPB1, HLA-DQA1, HLA-DQA2, HLA-DRA, HLA-DRB1, HLA-DRB5, HLA-E, RALB, RRAS2 |
| Cardiac Hypertrophy Signaling (Enhanced) | 2,83 | -1,279 | ADCY1, ADCY2, CD40LG, CSF2, FASLG, IGF1, IL12RB2, IL15RA, IL18R1, IL18RAP, IL1R2, IL2RA, IL4R, IL5, JAK2, MAP3K8, PDE4A, PDE7B, PRKCE, RALB, RRAS2, RYR1, TGFBR3, TNFSF10, TNFSF13B, WNT5B |
| Regulation of Cellular Mechanics by Calpain Protease | 1,78 | -1 | CAPN2, CCNA1, CDK6, RALB, RRAS2, VCL |
| STAT3 Pathway | 3,43 | -0,816 | IGF1, IL12RB2, IL15RA, IL18R1, IL18RAP, IL1R2, IL2RA, IL4R, JAK2, RALB, RRAS2, TGFBR3 |
| Sphingosine-1-phosphate Signaling | 1,36 | -0,816 | ACER2, ADCY1, ADCY2, CASP10, CASP4, PDGFD, S1PR5 |
| Gαi Signaling | 1,23 | -0,816 | ADCY1, ADCY2, CCR4, GNG4, RALB, RRAS2, TBXA2R |
| Coronavirus Pathogenesis Pathway | 1,24 | -0,707 | ADAM17, DDX58, FASLG, IRF7, STAT1, STAT2, STING1, TRIM25 |
| Calcium-induced T Lymphocyte Apoptosis | 3,34 | -0,707 | CAPN2, CD4, HLA-DOB, HLA-DQA1, HLA-DRA, HLA-DRB1, HLA-DRB5, PRKCE |
| Coronavirus Pathogenesis Pathway | 1,24 | -0,707 | ADAM17, DDX58, FASLG, IRF7, STAT1, STAT2, STING1, TRIM25 |
| Dendritic Cell Maturation | 2,73 | -0,577 | CD40LG, COL1A1, CSF2, HLA-DOB, HLA-DQA1, HLA-DRA, HLA-DRB1, HLA-DRB5, HLA-E, JAK2, MYD88, STAT1, STAT2 |
| Macropinocytosis Signaling | 1,73 | -0,447 | CSF1, MET, PDGFD, PRKCE, RALB, RRAS2 |
| GM-CSF Signaling | 1,35 | -0,447 | CSF2, JAK2, RALB, RRAS2, STAT1 |
| IL-15 Signaling | 1,25 | -0,447 | CSF2, IL15RA, JAK2, RALB, RRAS2 |
| NF-κB Activation by Viruses | 1,23 | -0,447 | CD4, EIF2AK2, PRKCE, RALB, RRAS2 |
| PPARα/RXRα Activation | 1,4 | -0,333 | ADCY1, ADCY2, FASN, HELZ2, IL18RAP, IL1R2, JAK2, RALB, RRAS2, TGFBR3 |
| Th1 Pathway | 7,4 | -0,302 | CD4, CD40LG, CD8A, HLA-DOB, HLA-DPA1, HLA-DPB1, HLA-DQA1, HLA-DQA2, HLA-DRA, HLA-DRB1, HLA-DRB5, IL12RB2, IL18R1, JAK2, KLRC1, LGALS9, STAT1 |

### Top canonical pathways revealed that interferons regulated signaling in DCs from HIV conditioned DC-T cell cocultures

Several top pathways of DCs from the mock and HIV treated cocultures were next examined as transcriptome heatmap profiles displaying all donors. The Interferon signaling pathway showed upregulation of JAK2, IFI36 and IFITM (**Figure 6A**). In addition, the pathway Activation of IRF by Cytosolic PRRs showed upregulation of ADAR, STAT2 and IRF7 (**Figure 6B**), and the pathway Pattern recognition receptors showed increased expression of FASLG, TNFS10 and MYD88 (**Figure 6C**). A clear difference in HIV versus mock exposed DCs was the activation of the Retinoic Acid Mediated Apoptosis Signaling pathway (**Figure 6D**) in which an increase in the signaling molecules PARP11, PARP10, and PARP12 were seen in the HIV exposed DC groups. Next, we assessed top upstream regulators and factors predicted to be activated including IFNa2, IFNL1, IFNβ, IFNγ, and the transcriptional regulators IRF7, STAT1, and IRF1 (**Table 3**). Top upstream regulators predicted to be inhibited were MAPK1, PNPT1, and IL1RN and the transcriptional regulators NKX2-3, TRIM24, and ETV6-RUNX1 (**Table 4**). When performing an analysis focusing on inflammation and infection associated factors in IPA, we found a strong Z-score for IFNβ and IFNA2 signaling as well as for IRF1 and IRF7 signaling (**Figure 6E**). Taken together these data suggest a strong activation of IFN signaling that might play a role in the induction of tolerogenic DCs and impaired T cell proliferation in the HIV DC-T cell coculture.

**Figure 6.**
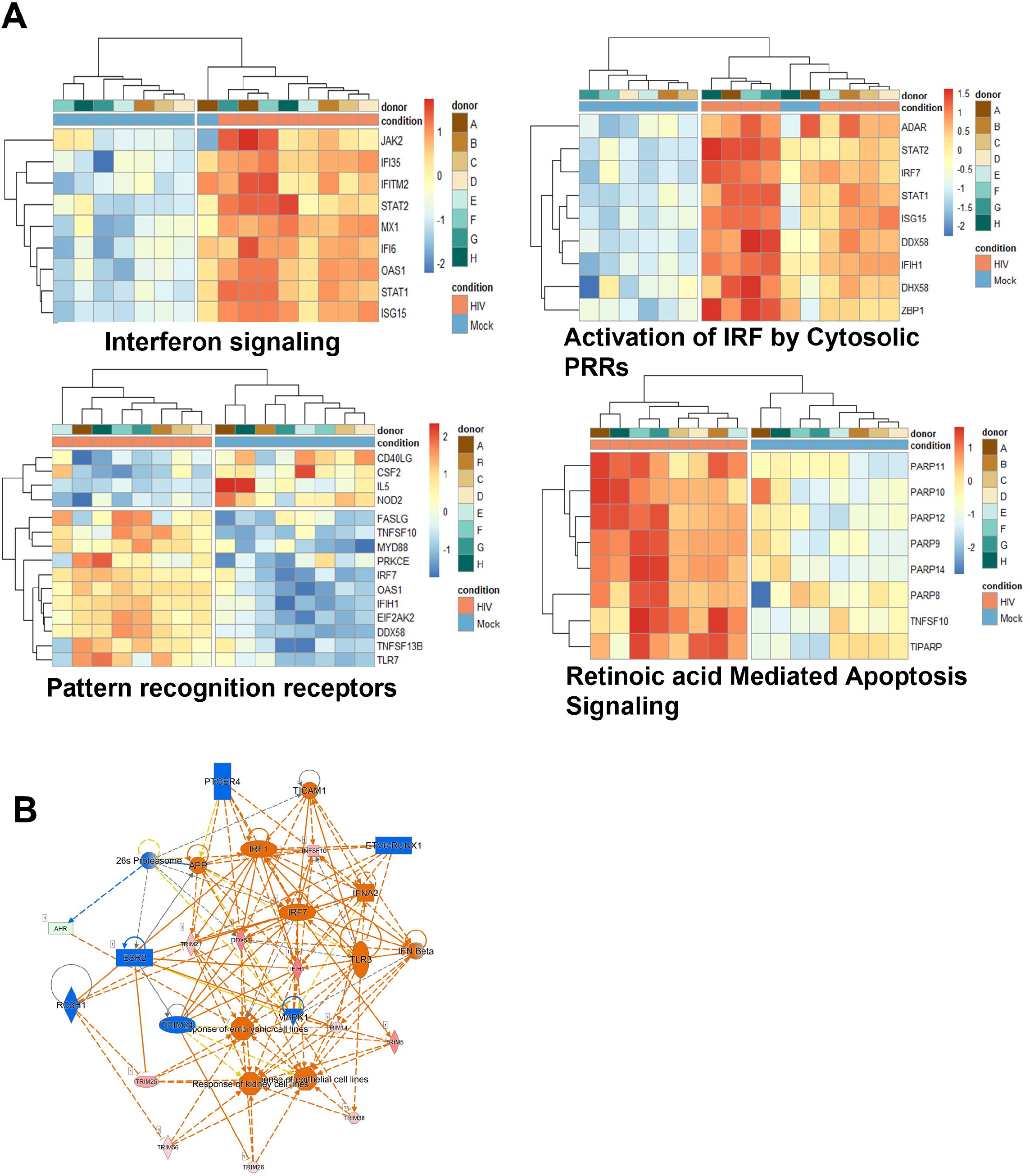
The top infection, inflammation, and immunology pathway regulators in HIV exposed DC cocultures. Transcriptomic data set including data from 8 individual donors/experiments was analyzed using pheatmap package to provide hierarchic clustered heatmaps of Interferon signaling **(A)**, Activation by Cytosolic PRRs **(B)**, Pattern recognition receptors **(C)** and Retinoic Acid Mediated Apoptosis Signaling **(D)** pathways with genes recognized in IPA to belong to these pathways. Transcriptomic data set from 8 individual donors/experiments was analyzed for Pathway regulators limited to infection, inflammation and immunology and were visualized as network with a threshold for p-values set to -log 1.3 (p<0.05) and presented as positive activation Z-score in orange and negative in blue **(E)**.

**Table 3:**
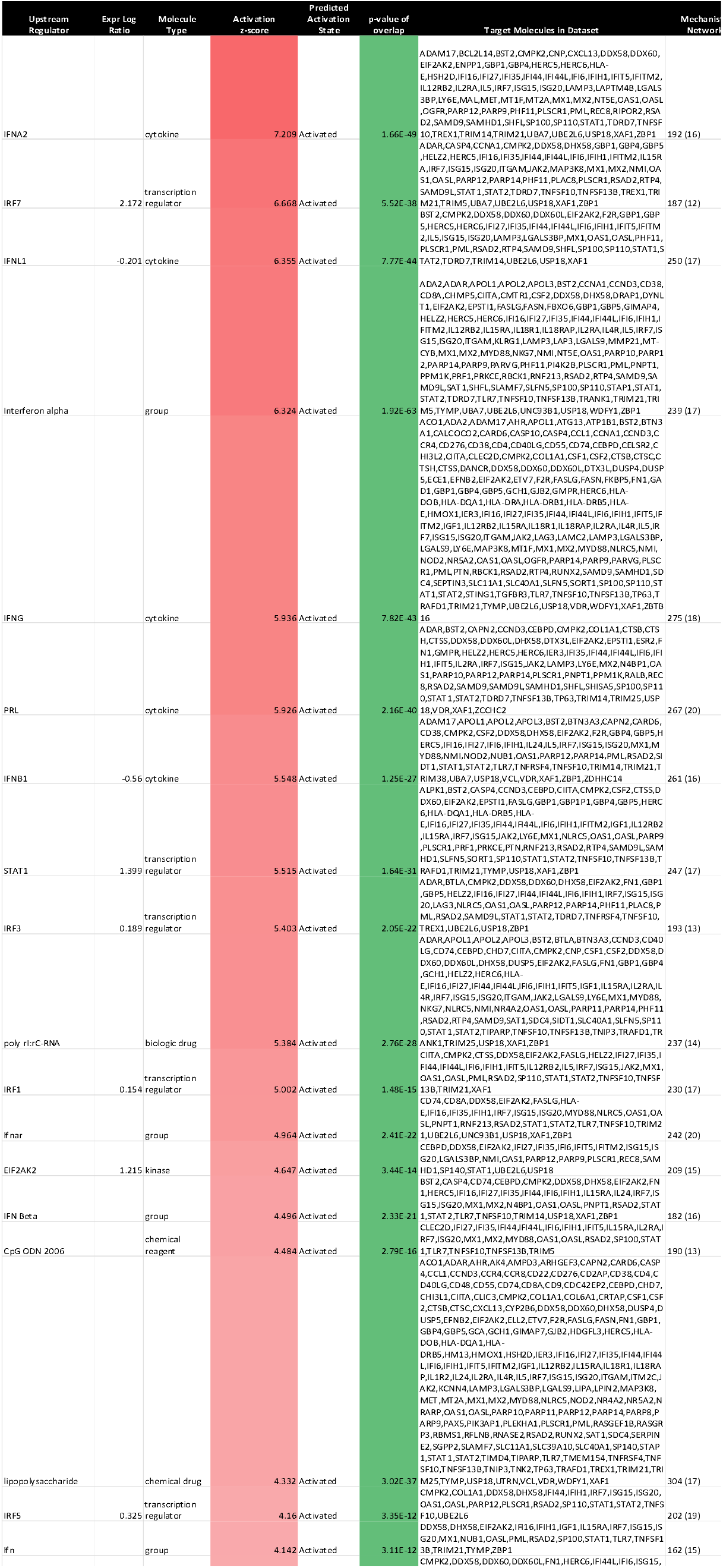
Upstream regulators with Positive Z-score

**Table 4:**
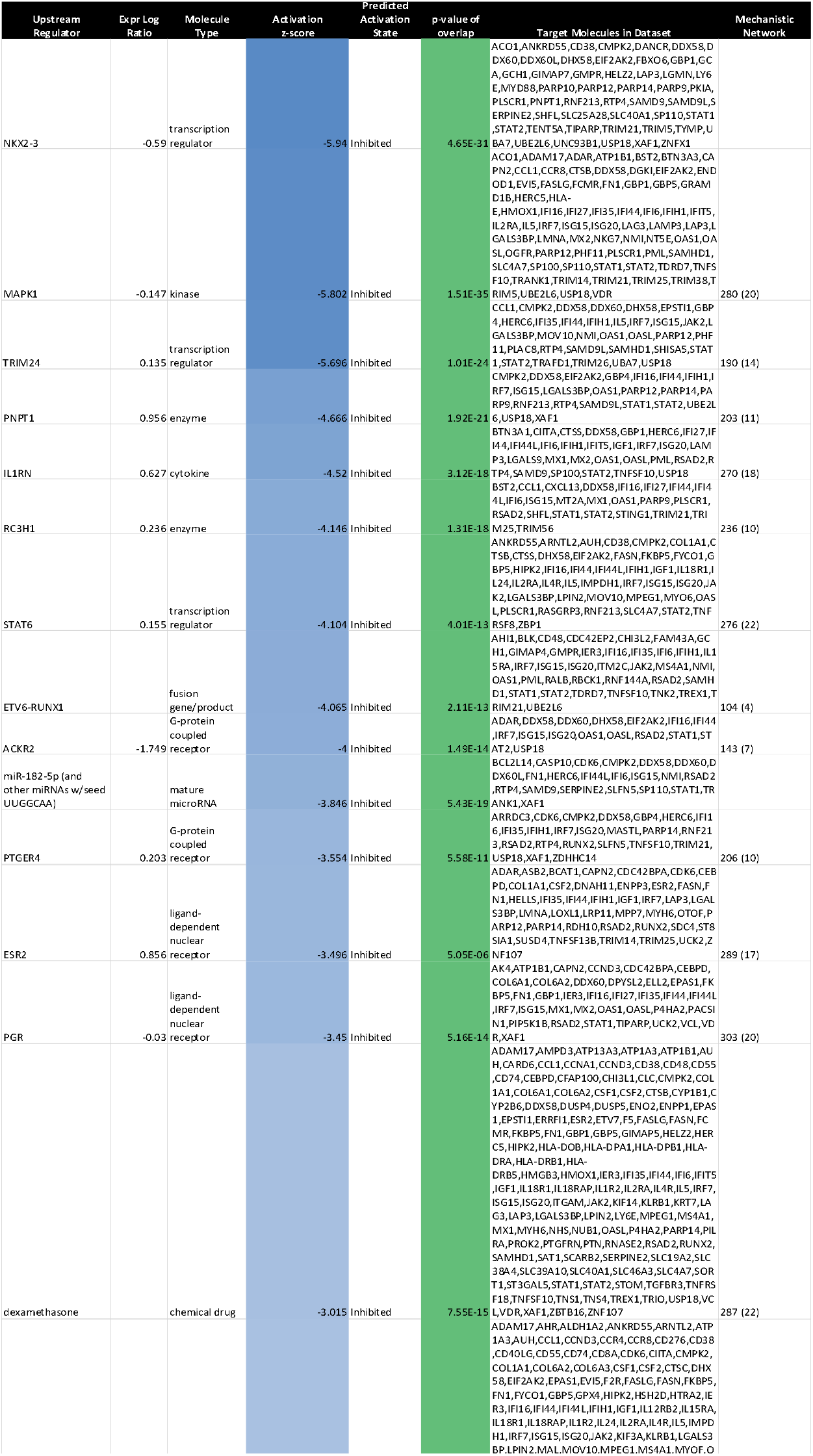
Upstream regulators with Negative Z-score

### DCs from HIV conditioned DC-T cell cocultures have altered T cell activating profile

The transcriptome data showed that T cell exhaustion signaling pathway was among the top pathways with a positive Z-score activated in the HIV exposed DCs from the DC-T cell cocultures (**Table 1**). The TH1 and TH2 Activation pathway and the Antigen presentation pathway were the top two canonical pathways affected in the DCs isolated from the HIV DC-T cell coculture but without clear activation or inhibition pattern (**Supplementary Table 1**). When dissecting the individual donor profiles for DCs isolated from the mock and HIV treated DC-T cell cocultures, we found that there were increased Z-scores for IL12RB2, LAG3 and JAK2, factors that have been associated with T cell exhaustion (**Figure 7A**). The analysis also showed an increase in TH1 signaling (**Figure 7B**) and TH2 signaling (**Figure 7C**) with positive Z-scores for IL12RB2, KLRC1, JAK2 and STAT1. The STAT3 pathway was also affected (**Figure 7D**) with positive Z-scores for IL15RA, JAK2 and IL18R1.

**Figure 7.**
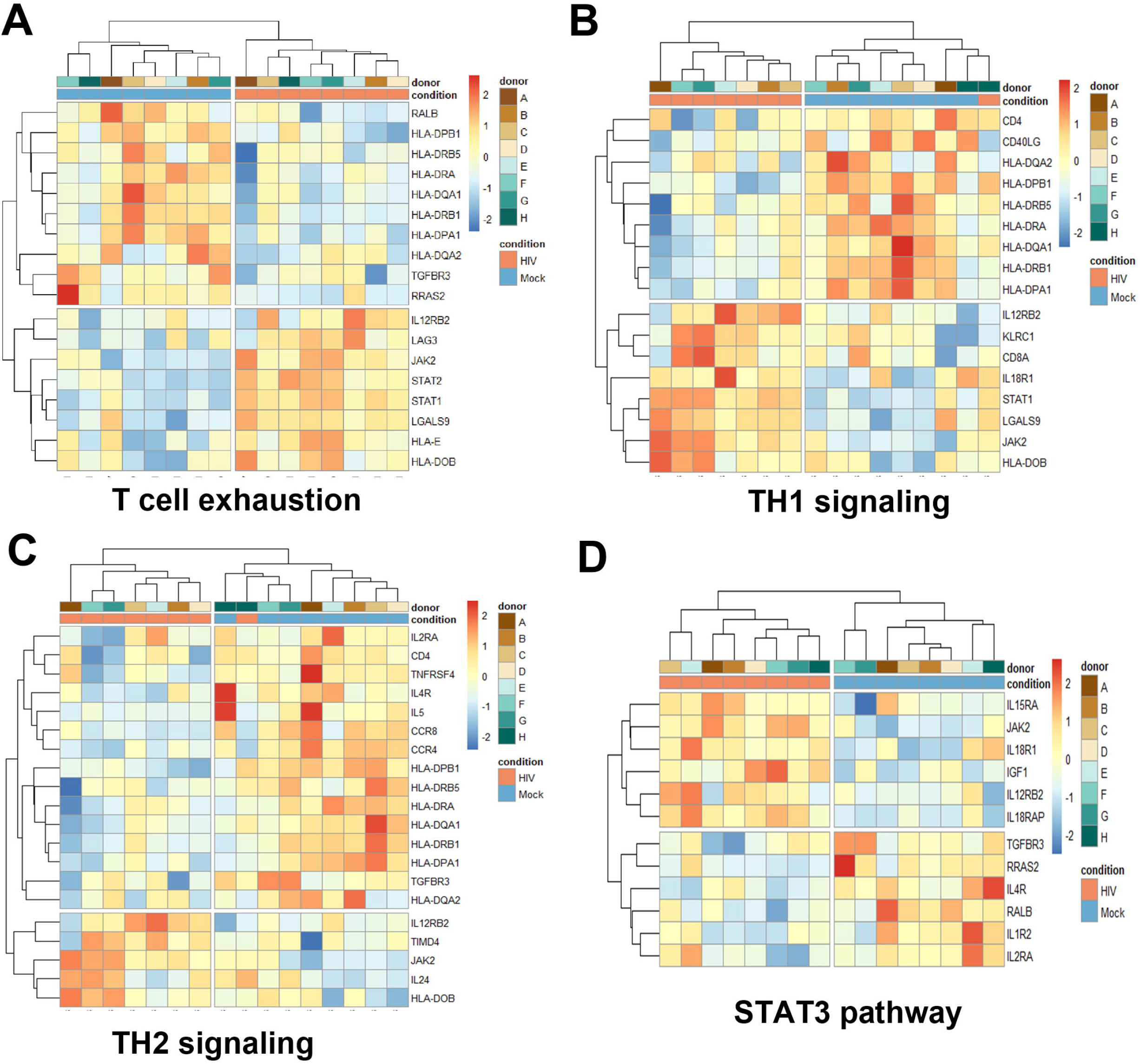
Transcriptomic data shows an activation of molecules involved in T cell exhaustion. Transcriptomic data set with data from 8 individual donors/experiments was analyzed using pheatmap package to provide hierarchic clustered heatmaps of Th1 signaling **(A)**, Th2 signaling **(B)**, STAT3 pathway **(C)**, and ICOS-ICOSL Signaling in T Helper Cells **(D)** pathways with genes recognized in IPA to belong to these pathways.

### Transcriptomic profiles of HIV exposed mature DCs after DC-T cell interaction had a high match score with different data sets of type I IFNs polarized conditions

We next performed match analysis to examine the match between our transcriptomic data of HIV exposed DCs of T cell cocultures with published data sets of human transcriptomic studies (**Figure 8A**). We set the cut off for the data sets with a minimum of 86% Canonical pathway Z-score match and minimum 76% Z-score overall score match. The top data sets included conditions characterized by type I IFN signaling; Influenza infection, Yellow fever vaccine, Sjogren’s syndrome, IFNβ treatment, influenza A infection of Lung adenocarcinoma, hepatocellular carcinoma treated with IFNα2a, and IFNβ treatment of multiple sclerosis (**Supplementary table 2**). Among the top five canonical pathways four were shared in the data sets; the Coronavirus pathogenesis pathway, The interferon signaling, Systemic Lupus Erythematosus in B cell signaling pathway, and neuroinflammation signaling pathway, whereas Role of hypercytokinemia/hyperchemokinemia in the pathogenesis of Influenza was not enriched in our data set (**Figure 8A**). The following inhibited pathways in our data sets stood out ICOS-IOCSL signaling in T helper cells, HMGB1 signaling, Role of NFAT in regulation of the immune responses, Dendritic cell maturation, and TH1 pathway. A pathway that was activated in our set but not in the other sets was the PD1-PDL1 cancer immunotherapy pathway. Analysis of the Upstream regulators showed an activated Z-score for an array of type IFN signaling molecules including IFNG, IFNA, IFNA2 and IRF7 for our and all matching data sets (**Figure 8B**). Comparing the different conditions, it was evident that type I IFN signaling was activated and regulators of inflammation including molecules such as NKX2-3, MAPK1, TRIM24, STAT6, IL1RN, and SOCS1 were suppressed. IRGM and IRM1 were inhibited in the other data sets but not in ours, whereas there was lower activation of IL-2, IL6, and NFkB (complex) in our data set, compared to the other data sets (**Figure 8B**).

**Figure 8.**
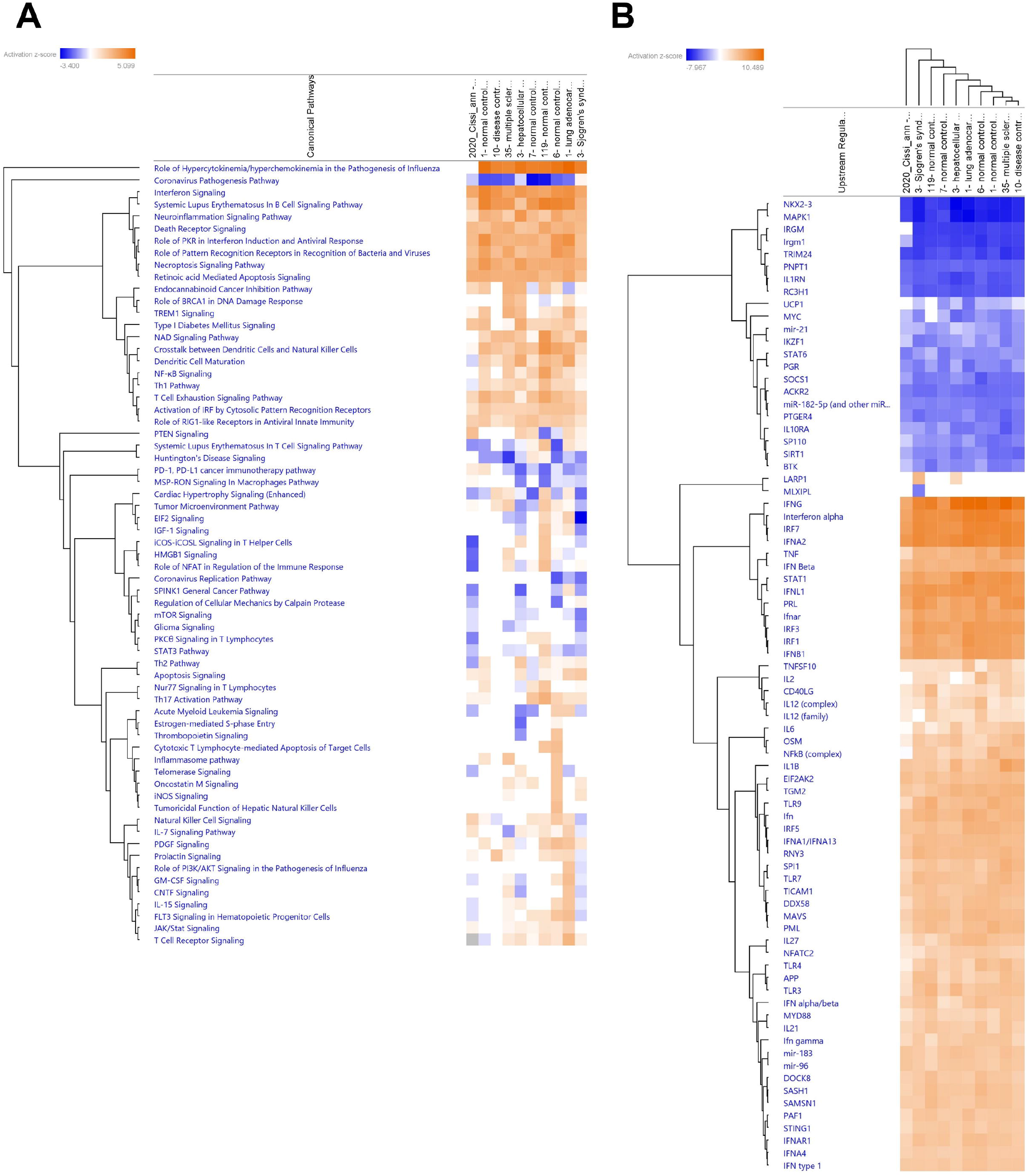
The HIV induced tolerogenic DCs showed in an analysis match a high overlap with type I IFN exposure, influenza infection and Sjogren syndrome data. Transcriptomic tolerogenic DC data set from 8 individual donors/experiments was analyzed using a match analysis with min 86% match in IPA and the matching data sets were 1- lung adenocarcinoma (LUAD) [alveoli] Infection_influenza A 12505, 10- normal control [peripheral blood] NA 8426, 31- multiple sclerosis (MS) [peripheral blood] NA 3961, 3- hepatocellular carcinoma (LIHC) [liver] IFN alpha 2a 14827, 3- Sjogren’s syndrome (SS) [peripheral blood] NA 16562, 5- relapsing-remitting MS (RRMS) [peripheral blood] NA 13564, 1- asthma [nasal epithelium] IFN alpha 8986, 7- normal control [peripheral blood] Infection_influenza A 6126, and 10- disease control [peripheral blood] NA 3340). Transcriptomic data set match analysis with top canonical pathways shown as hierarchic clustered heatmaps with a threshold for p-values set to -log 1.3 (p<0.05) and Z-score 1.5 **(A)**. Transcriptomic data set match analysis with Upstream regulator shown as hierarchic clustered heatmaps with a threshold for p-values set to log 1.3 (p<0.05) and Z-score 4 **(B)**.

## DISCUSSION

The interaction between immune cells and the network of signaling cascades that the cellular crosstalk induce are highly complex, and the immune response is dependent on cumulative signaling by all the co-stimulatory and co-inhibitory receptors and ligands. DCs simultaneously express multiple activating and inhibitory factors, and depending on the balance, can activate or suppress the ensuing T cell responses (Wilson et al. 2012). Our current investigations advance the understanding of the impaired immune activation that occurs when HIV is present during DC priming of T cells and the induction of suppressive T cells (Che et al. 2010, Che et al. 2012, Shankar et al. 2011). We found that mere exposure of DCs to HIV had no significant effect on the DCs’ gene expression levels of the co-inhibitory factors, and is in accordance with previous findings, including our own, showing very little or almost no effect of HIV exposure alone on the DC phenotype (Lubong Sabado et al. 2009, Manel et al. 2010, Hanley et al. 2010). Instead, it was the contact with the suppressive T cells that induced the alteration of the mature DCs phenotype, expressing an array of co-inhibitory molecules including PDL1, HVEM, Gal-9, B7H3, PDL2, CD85, and CD30L, as mere HIV exposure alone did not alter their phenotype. These findings suggest that DC-T cell cellular interaction could lead to a bidirectional expression of tolerogenic molecules. While there exists a wealth of information regarding the immunosuppressive markers on immune cells, there is a lack of understanding of the mechanisms that induce and direct immune suppression. In our settings, T cell interaction induces strong and long-lasting type I IFN responses in the DCs, compared to the transient type I IFN activation seen after HIV exposure of DCs in the absence of T cells, and sustained type I IFN signaling could be one factor that triggers the shift to a tolerogenic DC phenotype and the subsequent induction of an immunosuppressive environment.

Since then, a large number of immunomodulatory molecules and pathways involved in controlling T cell response activation has been defined (Chen and Flies 2013). Among the first costimulatory molecules defined to play an important role in T cell activation was the CD80 and CD86, which both bind CD28 (Schonrich and Raftery 2019). Today, a lot is known about the results the engagement of co-inhibitory molecules such as PD1 and CTLA4 have on T cells. However, the reverse impact, i.e., the effect on antigen presenting cells (APCs) such as DCs, is less studied. PDL1 is among the most prominent negative immune checkpoint molecules expressed on the DC’s surface, and the PD1 axis is a potent suppressive pathway (Schonrich and Raftery 2019). The PD1-PDL1 interaction will send inhibitory signals into the T cells via PD1 and into the DCs via PDL1. Studies have shown that PDL1 can also interact with CD80 and contribute to T cell suppression (Butte et al. 2007, Fehervari 2019). PDL1 is upregulated in tumor cells, and immune cells such as APCs, including DCs, by e.g., IFNγ, IL-10, IFNα, and IFNβ (Bazhin et al. 2018, Garcia-Diaz et al. 2019, Morimoto et al. 2018, Curiel et al. 2003). The role of PDL1 appears to be more complex than to suppress the T cell responses against tumor cells, since its presence also can protect against the immune suppressive effect of sustained type I and II IFNs responses (Azuma et al. 2008). The consequence of receptor-ligand interaction on the cell function is complex and context dependent. Surprisingly, engagement of the T cell costimulatory molecule CD28 can under some circumstances decrease T cell proliferation (Silberman et al. 2012).

The increased expression of DcR2 on DCs present in HIV exposed cocultures is important since these decoy receptors reportedly bind to TRAIL and can facilitate T cell inhibition (Tawfik et al. 2019). We and others had previously shown that HIV infection could promote increased expression of TRAIL *in vitro* (Che et al. 2010) and *in vivo* (Herbeuval et al. 2005). Together, these results indicate a role of TRAIL-DcR2 signaling in HIV-associated immune suppression as seen herein from our *in vitro* observations. We found an elevated Gal-9 expression in the HIV exposed DCs after interaction with T cells in the coculture, and this β-galactosidase-binding lectin has been shown to promote an immune suppressive microenvironment with impaired T cell responses in different settings including HIV infection (Shahbaz et al. 2020). Gal-9 is increased during acute HIV infection and the levels remain elevated also in individuals controlling the infection (Tandon et al. 2014).

DCs orchestrate the fate of T cell responses and can efficiently regulate cellular effector functions through diverse mechanisms that range from synthesis of factors exerting broadly attenuating effects to the induction of antigen-specific T cell responses that result in immune tolerance. The mature DCs prime naïve T cells and drive their polarization toward different T helper and cytotoxic T cell subtypes. The DCs are affected by the local microenvironment that also will impact the type of T cell response induced. The transcriptome profile of DCs from the HIV DC-T cell cocultures showed an inhibition of the TH2 pathway with decreased expression of IL5, TNFRSF4 (OX40/CD134), CCR4, and CCR8 (Iberg and Hawiger 2020, Steinman, Hawiger and Nussenzweig 2003). The TH1 pathway was less affected but still downregulated, suggesting a more general immune suppression not limited to TH2. Increased activation of T cell exhaustion pathways also is in support of an HIV associated general immune suppression.

Many types of epithelial and immune cells are impaired in chronic diseases such as HIV. The local microenvironment, characterized by multiple cellular interactions, bi-directional cell signaling and the release of inflammatory mediators, play a major role in the development of impaired immune function (Wang et al. 2019, Plaeger et al. 2012, Mehraj et al. 2014). There is a paucity in the understanding of the mechanisms leading to an immune suppressive microenvironment. It is clear from our study that the type I IFN-induced responses are stronger and last longer in HIV exposed DCs that have been in contact with T cells compared to the transient type I IFN activation seen when DCs have been exposed to HIV but not cocultured with T cells. It has been suggested that during chronic conditions such as viral persistence the type I IFN can have suppressive effects on the immune responses (Wilson et al. 2013). In addition, type I IFNs can induce tolerogenic/suppressive DCs in LCMV infection, HIV infection, tuberculosis, and cancer (Cunningham et al. 2016). Furthermore, the expression of PDL1 and the immune suppressive IL10, were highly dependent on type I IFN signaling in persistent viral infections (Teijaro et al. 2013). It is becoming clear from several studies, that for type I IFNs, the magnitude and timing of expression are essential for the outcome of the infection and quality of the immune responses (Cunningham et al. 2016, Wilson et al. 2013, King and Sprent 2021). The antiviral type I IFN response and the inflammatory response are balanced by mutual crosstalk (McNab et al. 2015). The level of inflammation in the DCs and in our HIV coculture system is in general low and this could be the result of an imbalance reflected by a strengthened type I IFN responses in these systems. The role of type I IFNs in HIV pathogenesis has been debated. Early in HIV infection, increased IFNα levels are considered to restrict infection, i.e., beneficial, whereas elevated levels later in disease are associated with disease progression with increased viral loads and decreased CD4 T cell counts (Hardy et al. 2013, Stacey et al. 2009). Furthermore, it has been suggested that there is a link between type I IFN and pathogenic disease and progression to AIDS in SIV (Jacquelin et al. 2009, Bosinger and Utay 2015). These findings in HIV infected individual and SIV experimental models support a role of type I IFNs in the development of impaired immune responses and the induction of suppressive/tolerogenic DCs in our study.

DCs can also modulate T cell differentiation by modifying metabolic parameters in the T cell microenvironment by production and release of retinoic acid (Sun et al. 2007). The transcriptomic data showed activation of the retinoic acid mediated apoptosis signaling pathway. In addition, STAT3, and STAT1 and STAT2 can be activated by type I IFNs and STAT3 is known to confer immunosuppressive functions (Yu, Kortylewski and Pardoll 2007). We identified an upregulation of STAT3 signaling in transcriptome profiles of the HIV exposed DCs after T cell interaction, and we know from our previous findings that STAT3 blockade in our ex vivo model of DC-T cell coculture can restore the DC T cell priming capacity after HIV exposure (Che et al. 2012).

Taken together, this study emphasizes that exposure of mature DCs to HIV significantly altered the overall balance in the expression of immune checkpoint molecules in favor of co-inhibitory molecules after the interaction with T cells. These findings, shed a new light on the understanding of HIV that potentially skews DCs to differentiate into tolerogenic/suppressive DC phenotype where cellular contact with the T cells alters their functionality in which the IFN type I signaling could play an integral role. Given the emergence of a wider network of co-inhibitory molecules in HIV infection, it is important now to investigate the specific role this environment plays in HIV pathogenesis in infected individuals.

## Acknowledgments

We thank the Biological Products Core of the AIDS and Cancer Virus Program, SAIC Frederick, Inc., National Cancer Institute, Frederick, MD, USA for generously providing HIV. The data handling was enabled by resources provided by the Swedish National Infrastructure for Computing (SNIC) at UPPMAX partially funded by the Swedish Research Council through grant agreement no. 2018-05973.

