## Supplementary figures for "HIV-1 Induction of Tolerogenic DCs is Mediated by Cellular Interaction with Suppressive T Cells"

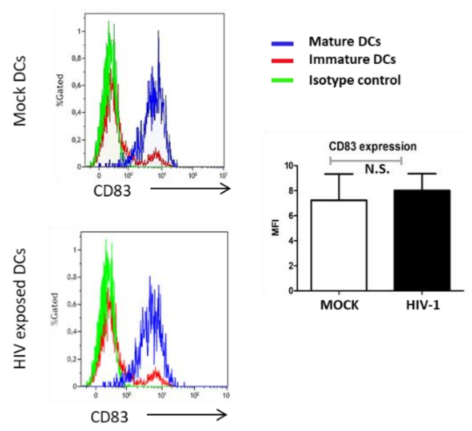


**Supplementary Figure 1. HIV-1 does not alter the DCs CD83 expression/maturation status compared to mock.** Immature and mature DCs were exposed to HIV for 24h and stained with anti-CD83 antibody or isotype control antibody and analysed by flow cytometry and visualized by representative histograms (left) and MFI graph (right). N =3.

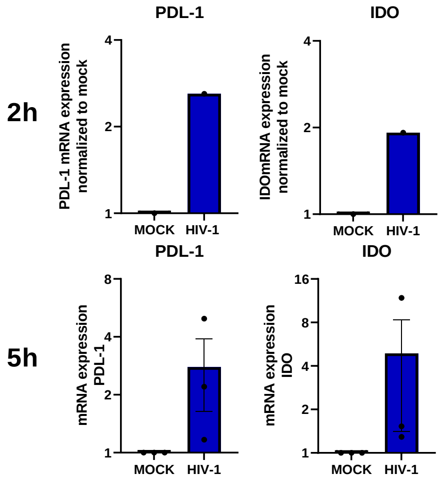


**Supplementary Figure 2. Upregulation of negative costimulatory molecule and modulatory factors occurs fast in DCs after contact the Day 7 DC-T cell coculture.**

mRNA Expression of PDL-1 and IDO after 2h and 5h DC re-exposure of co-cultures was investigated with qPCR. Expression is shown as relative expression compared with its respective unexposed (mock) control which are stated as 1. Results are shown as mean with standard deviation. N =1-3.

### Supplementary Table 1. Top Canonical pathways

| Canonical Pathways | -log(p-value) | z-score | Molecules |
|---|---|---|---|
| **Th1 and Th2 Activation Pathway** | 11.8 | NaN | CCR4,CCR8,CD4,CD40LG,CD8A,HLA-DOB,HLA-DPA1,HLA-DPB1,HLA-DQA1,HLA-DQA2,HLA-DRA,HLA-DRB1,HLA-DRB5,IL12RB2,IL18R1,IL24,IL2RA,IL4R,IL5,JAK2,KLRC1,LGALS9,STAT1,TGFBR3,TIMD4,TNFRSF4 |
| **Antigen Presentation Pathway** | 9.45 | NaN | CD74,CIITA,HLA-DOB,HLA-DPA1,HLA-DPB1,HLA-DQA1,HLA-DQA2,HLA-DRA,HLA-DRB1,HLA-DRB5,HLA-E,NLRC5 |
| **Th2 Pathway** | 8.95 | -1.265 | CCR4,CCR8,CD4,HLA-DOB,HLA-DPA1,HLA-DPB1,HLA-DQA1,HLA-DQA2,HLA-DRA,HLA-DRB1,HLA-DRB5,IL12RB2,IL24,IL2RA,IL4R,IL5,JAK2,TGFBR3,TIMD4,TNFRSF4 |
| **Th1 Pathway** | 7.4 | -0.302 | CD4,CD40LG,CD8A,HLA-DOB,HLA-DPA1,HLA-DPB1,HLA-DQA1,HLA-DQA2,HLA-DRA,HLA-DRB1,HLA-DRB5,IL12RB2,IL18R1,JAK2,KLRC1,LGALS9,STAT1 |
| **T Cell Exhaustion Signaling Pathway** | 6.38 | 1.414 | BTLA,HLA-DOB,HLA-DPA1,HLA-DPB1,HLA-DQA1,HLA-DQA2,HLA-DRA,HLA-DRB1,HLA-DRB5,HLA-E,IL12RB2,JAK2,LAG3,LGALS9,RALB,RRAS2,STAT1,STAT2,TGFBR3 |
| **Interferon Signaling** | 6.37 | 3 | IFI35,IFI6,IFITM2,ISG15,JAK2,MX1,OAS1,STAT1,STAT2 |
| **MSP-RON Signaling In Macrophages Pathway** | 6.28 | 0.258 | CIITA,GAB2,HLA-DOB,HLA-DPA1,HLA-DPB1,HLA-DQA1,HLA-DQA2,HLA-DRA,HLA-DRB1,HLA-DRB5,ITGAM,JAK2,RALB,RRAS2,STAT1 |
| **Allograft Rejection Signaling** | 6.18 | NaN | CD40LG,FASLG,HLA-DOB,HLA-DPA1,HLA-DPB1,HLA-DQA1,HLA-DQA2,HLA-DRA,HLA-DRB1,HLA-DRB5,HLA-E,IL5,PRF1 |
| **T Helper Cell Differentiation** | 6.15 | NaN | CD40LG,HLA-DOB,HLA-DQA1,HLA-DRA,HLA-DRB1,HLA-DRB5,IL12RB2,IL18R1,IL2RA,IL4R,IL5,STAT1 |
| **Autoimmune Thyroid Disease Signaling** | 6.12 | NaN | CD40LG,FASLG,HLA-DOB,HLA-DQA1,HLA-DRA,HLA-DRB1,HLA-DRB5,HLA-E,IL5,PRF1 |
| **Systemic Lupus Erythematosus In B Cell Signaling Pathway** | 5.57 | 1.877 | CCND3,CD22,CD40LG,CSF2,FASLG,IFIH1,IL5,IRF7,ISG15,ISG20,JAK2,MYD88,PIK3AP1,PRKCE,RALB,RASGRP3,RRAS2,STAT1,STAT2,STING1,TLR7,TNFSF10,TNFSF13B |
| **Altered T Cell and B Cell Signaling in Rheumatoid Arthritis** | 5.16 | NaN | CD40LG,CSF1,CSF2,CXCL13,FASLG,HLA-DOB,HLA-DQA1,HLA-DRA,HLA-DRB1,HLA-DRB5,TLR7,TNFSF13B |
| **Death Receptor Signaling** | 5.06 | 2.309 | CASP10,FASLG,HTRA2,LMNA,PARP10,PARP11,PARP12,PARP14,PARP8,PARP9,TIPARP,TNFSF10 |
| **Hepatic Fibrosis / Hepatic Stellate Cell Activation** | 4.76 | NaN | CD40LG,COL1A1,COL6A1,COL6A2,COL6A3,CSF1,ECE1,FASLG,FN1,IGF1,IL18RAP,IL1R2,IL4R,MET,MYH6,PDGFD,STAT1 |
| **Role of Pattern Recognition Receptors in Recognition of Bacteria and Viruses** | 4.6 | 2.121 | CD40LG,CSF2,DDX58,EIF2AK2,FASLG,IFIH1,IL5,IRF7,MYD88,NOD2,OAS1,PRKCE,TLR7,TNFSF10,TNFSF13B |
| **Crosstalk between Dendritic Cells and Natural Killer Cells** | 4.47 | 0.302 | CD40LG,CSF2,FASLG,HLA-DRA,HLA-DRB1,HLA-DRB5,HLA-E,IL15RA,PRF1,TLR7,TNFSF10 |
| **OX40 Signaling Pathway** | 4.42 | NaN | CD4,HLA-DOB,HLA-DPA1,HLA-DPB1,HLA-DQA1,HLA-DQA2,HLA-DRA,HLA-DRB1,HLA-DRB5,HLA-E,TNFRSF4 |
| **Graft-versus-Host Disease Signaling** | 4.34 | NaN | FASLG,HLA-DOB,HLA-DQA1,HLA-DRA,HLA-DRB1,HLA-DRB5,HLA-E,PRF1 |
| **Activation of IRF by Cytosolic Pattern Recognition Receptors** | 4.26 | 1 | ADAR,DDX58,DHX58,IFIH1,IRF7,ISG15,STAT1,STAT2,ZBP1 |
| **Type I Diabetes Mellitus Signaling** | 4.24 | 2 | FASLG,GAD1,HLA-DOB,HLA-DQA1,HLA-DRA,HLA-DRB1,HLA-DRB5,HLA-E,JAK2,MYD88,PRF1,STAT1 |
| **Communication between Innate and Adaptive Immune Cells** | 4.16 | NaN | CD4,CD40LG,CD8A,CSF2,HLA-DRA,HLA-DRB1,HLA-DRB5,HLA-E,IL5,TLR7,TNFSF13B |
| **Oncostatin M Signaling** | 3.79 | 0 | CHI3L1,EPAS1,JAK2,MT2A,RALB,RRAS2,STAT1 |
| **PD-1, PD-L1 cancer immunotherapy pathway** | 3.77 | 0.707 | HLA-DOB,HLA-DPA1,HLA-DPB1,HLA-DQA1,HLA-DQA2,HLA-DRA,HLA-DRB1,HLA-DRB5,HLA-E,IL2RA,JAK2 |
| **Retinoic acid Mediated Apoptosis Signaling** | 3.63 | 2.828 | PARP10,PARP11,PARP12,PARP14,PARP8,PARP9,TIPARP,TNFSF10 |
| **STAT3 Pathway** | 3.43 | -0.816 | IGF1,IL12RB2,IL15RA,IL18R1,IL18RAP,IL1R2,IL2RA,IL4R,JAK2,RALB,RRAS2,TGFBR3 |

### Supplementary Table 2. Analysis Match with other data sets

| Analysis Name | case. diseasestate | case. tissue | case. treatment | comparison category | comparison contrast | comparison id | weblink | CP z-score | UR z-score | CN z-score | DE z-score | z-score overall score |
|---|---|---|---|---|---|---|---|---|---|---|---|---|
| 1- lung adenocarcinoma (LUAD) [alveoli] Infection_influenza A 12505 | lung adenocarcinoma (LUAD) | alveoli | Infection_influenza A | Treatment vs. Control | Infection => influenza A vs mock | GSE31474.GPL570.test1 | https://www.ncbi.nlm.nih.gov/geo/query/acc.cgi?acc=GSE31474 | 86.6 | 80.62 | 72.11 | 72.89 | 78.06 |
| 10- normal control [peripheral blood] NA 8426 | normal control | peripheral blood | NA | Treatment1 vs. Treatment2 | Vaccine:SamplingTime => YF-VAX -> day 7 vs day 3 | GSE13699.GPL6104.test10 | https://www.ncbi.nlm.nih.gov/geo/query/acc.cgi?acc=GSE13699 | 86.6 | 82.46 | 67.08 | 75 | 77.79 |
| 31- multiple sclerosis (MS) [peripheral blood] NA 3961 | multiple sclerosis (MS) | peripheral blood | NA | Treatment vs. Control | SamplingTime:CellType => helper T cell -> 24 hours after the first treatment vs baseline | GSE60424.GPL15456.DESeq2.test31 | http://www.ncbi.nlm.nih.gov/geo/query/acc.cgi?acc=GSE60424 | 86.6 | 82.46 | 72.11 | 68.47 | 77.41 |
| 3- hepatocellular carcinoma (LIHC) [liver] IFN alpha 2a 14827 | hepatocellular carcinoma (LIHC) | liver | IFN alpha 2a | Treatment vs. Control | SamplingTime => 12 hours vs baseline | GSE48400.GPL10558.test3 | https://www.ncbi.nlm.nih.gov/geo/query/acc.cgi?acc=GSE48400 | 86.6 | 79.37 | 60 | 81.01 | 76.75 |
| 3- Sjogren's syndrome (SS) [peripheral blood] NA 16562 | Sjogren's syndrome (SS) | peripheral blood | NA | Disease vs. Normal | DiseaseState => Sjogren's syndrome (SS) vs normal control | GSE66795.GPL10558.test3 | http://www.ncbi.nlm.nih.gov/geo/query/acc.cgi?acc=GSE66795 | 86.6 | 81.85 | 71.41 | 66.14 | 76.5 |
| 5- relapsing-remitting MS (RRMS) [peripheral blood] NA 13564 | relapsing-remitting MS (RRMS) | peripheral blood | NA | Treatment vs. Control | DiseaseStage:TreatmentStatus => in remission -> IFN beta 1a vs none | GSE41890.GPL6244.test5 | http://www.ncbi.nlm.nih.gov/geo/query/acc.cgi?acc=GSE41890 | 79.06 | 83.67 | 70 | 72.89 | 76.4 |
| 1- asthma [nasal epithelium] IFN alpha 8986 | asthma | nasal epitheliu m | IFN alpha | Treatment vs. Control | Treatment => IFN alpha vs none | GSE19182.GPL6244.test1 | http://www.ncbi.nlm.nih.gov/geo/query/acc.cgi?acc=GSE19182 | 86.6 | 83.07 | 69.28 | 66.14 | 76.27 |
| 7- normal control [peripheral blood] Infection_influenza A 6126 | normal control | peripheral blood | Infection_influenza A | Treatment vs. Control | SamplingTime[hpi]:VirusStrain => 6 -> A/Kawasaki/UTK-4/2009(H1N1) vs NA | GSE100865.GPL16686.test7 | https://www.ncbi.nlm.nih.gov/geo/query/acc.cgi?acc=GSE100865 | 86.6 | 82.46 | 64.81 | 70.71 | 76.15 |
| 10- disease control [peripheral blood] NA 3340 | disease control | peripheral blood | NA | Treatment vs. Control | ExperimentGroup2 => anakinra before MMT with hypoglycemic event vs placebo before MMT with | GSE132781.GPL16791.DESeq2.test10 | https://www.ncbi.nlm.nih.gov/geo/query/acc.cgi?acc=GSE132781 | 86.6 | 81.24 | 65.57 | 70.71 | 76.03 |

Selected on Human disease data bas

CP = canonical pathway Z-Score, UR = upstream regulators Z-Score, DE = diseases and functions (downstream effect) Z-Score, CN = causal network Z-Score
